# First genomic resource of ‘*Candidatus* Phytoplasma pyri’ associated to pear

**DOI:** 10.1101/2025.01.23.634465

**Authors:** Florencia Ivette Alessio, Vanina Aylen Bongiorno, Oscar Hernan von Backzo, Luis Rogelio Conci, Franco Daniel Fernandez

## Abstract

This study reports the draft genome of *Candidatus* Phytoplasma pyri strain P1, isolated from Argentina, marking the first global sequencing of this species. The genome assembly consisted of 16 contigs, with a total length of 575,431 bp, a GC content of 20.35%, and 125X coverage. A total of 536 genes were annotated, including those related to metabolism, genetic information processing, and signaling. Phylogenetic analysis placed ‘*Candidatus* Phytoplasma pyri’ within the 16SrX group, supporting its classification as a distinct species from ‘*Candidatus* Phytoplasma mali’ and ‘*Candidatus* Phytoplasma prunorum’, despite their close evolutionary proximity. The investigation also identified 17 secreted proteins, including homologs of known effectors such as SAP39 and SAP41, as well as conserved proteins associated with virulence mechanisms. A homologue of the Imp immunodominant membrane protein was also characterized, revealing conserved gene organization across related strains. These findings contribute to the understanding of ‘*Candidatus* Phytoplasma pyri’ pathogenic and provide valuable genomic data for future research. This study highlights the importance of expanding phytoplasma genomics, as it provides a crucial first step in understanding ‘*Candidatus* Phytoplasma pyri’ and its role in plant pathology

## 1. Introduction

Phytoplasmas are obligate plant pathogens within the class *Mollicutes* that cause devastating diseases in a wide range of economically important crops worldwide, including fruit trees, vegetables, and ornamentals (Lee and Davis, 2000). These pathogens are primarily transmitted by phloem-feeding insect vectors like leafhoppers, planthoppers, and psyllids, contributing to their global distribution (Weintraub and Beanland, 2006). Due to their unculturable nature, phytoplasma identification relies heavily on molecular methods. The 16S rRNA gene is central to their classification, enabling the delineation of over 30 major groups through sequence analysis and RFLP (Lee et al., 1998). The classification of ‘*Candidatus* (Ca.) Phytoplasma’ species has traditionally been based on the 16S rRNA gene, with initial guidelines from 2004 (IRPCM, 2004) setting a 97.5% sequence similarity threshold for species delimitation, alongside criteria for distinct ecological or molecular characteristics. Updated guidelines in 2022 raised this threshold to 98.65% and incorporated analyses of housekeeping genes and whole-genome average nucleotide identity (ANI), while retaining the ecological differentiation criteria for closely related taxa (Bertaccini et al., 2022; Wei and Zhao, 2022). To date, approximately 50 ‘*Ca*. Phytoplasma’ species have been formally described (Bertaccini et al., 2022). Among these species, ‘*Ca*. Phytoplasma pyri’ (16SrX-C subgroup) is the causal agent of pear decline, a globally significant disease that severely impacts pear production, causing substantial damage and crop losses (Seemüller and Schneider, 2004). Infected pear trees display symptoms such as yellowing, reddening, loss of vigor, premature blooming, and, in some cases, total collapse and death of the tree (Sabaté et al., 2018). The progression of the disease and the symptoms exhibited are strongly influenced by both the rootstock and the disease stage (Seemüller et al., 2011).The significance of *‘Ca*. Phytoplasma pyri’ for pear production is steadily increasing, and it has been proposed for regulation as a quarantine organism in the European Union (A2 List) (EPPO Global Database, 2009). Argentina is the second-largest global producer of pears, with production concentrated in Río Negro, Neuquén, and Mendoza, covering 20,330 hectares. In 2022, the country produced 580,000 tons, with 71% exported, making pears its top fresh fruit export. Major markets include Brazil, the U.S., and Europe, underscoring Argentina’s importance in global fruit trade (Ministerio de Agricultura, Ganadería y Pesca, 2022). The first report of ‘*Ca*. Phytoplasma pyri’ in South America was made in Argentina, associated with peach trees exhibiting symptoms such as chlorotic leaves, ridges, and thickening of the central veins (Fernandez et al., 2017). Later that same year, its presence was detected in Chile in pears showing reddening symptoms (Facundo et al., 2017). In 2022, ‘*Ca*. Phytoplasma pyri’ was found in pear plants from the Río Negro production region in Argentina, exhibiting a marked symptom of reddening (Fernandez et al., 2019a). Given the lack of reference genomes for *Candidatus* Phytoplasma pyri globally, the present study aims to sequence and characterize the genome of a ‘*Ca*. Phytoplasma pyri’ isolate from Argentina.

## Materials and Methods

### 2.1 Plant sample and phytoplasma detection

Total DNA extraction (gDNA) was performed on a representative pear sample infected with phytoplasma (designated as P1) using the Monarch Genomic DNA Purification Kit (NEB, USA). The sample was collected from pear production fields located in General Roca (Rio Negro, Argentina 39°02′00″S 67°35′00″O). The extracted DNA was quantified using both a Nanodrop (N1000) and a Quantus Fluorometer (Promega). To confirm the presence of phytoplasma, gDNA was analyzed by PCR with universal primers P1/P7 (Deng & Hiruki, 1991; Schneider et al., 1995) for direct PCR, and R16F2n/R16R2 (Lee et al., 1993) for nested PCR. Amplicons were analyzed in agarose gel electrophoresis and stained with GelRed®. Identification of 16Sr group very accessed by RFLP analysis of PCR products (1.2 kb, using R16F2/R16R2 primers) were digested with *Mse*I, *Rsa*I, *Taq*I, and *Ssp*I enzymes (NEB, USA) following the manufacturer’s instructions. The restriction profiles were analyzed using agarose (1.5X:0.5X) gel electrophoresis, stained with GelRed® (Biotium, USA), and visualized using a UV transilluminator. The resulting patterns were compared with those described previously (Fernandez et al. 2019a).

### 2.2 Sequencing and assembling

For whole genome sequencing a hybrid approach was employed combining Illumina and ONT platforms. For Illumina sequencing, the library was processed on a NovaSeq using the TruSeq Nano DNA (350) kit. The reads underwent trimming using Trimmomatic v0.39 with parameters PE –phred33 ILLUMINACLIP LEADING:3 TRAILING:3 SLIDINGWINDOW:4:15 MINLEN:36. ONT sequencing involved library preparation with the ONT Ligation Sequencing Kit (SQK-LSK109) and sequencing on a MinION (FLO-MIN114). Basecalling was performed using Guppy v6.5.7 with the super high accuracy mode, setting a minimum q-score of 10. The assembly was performed using Unicycler version 0.5.0 (https://github.com/rrwick/Unicycler). Phytoplasma contigs were identified by conducting BLASTx searches against phytoplasma proteins local database. Raw reads were then mapped to these contigs using bowtie2 v2.3.5.1 and Minimap2 v2.15 in an iterative process until the assembly was complete. Final contigs were polished using two rounds of Polypolish (https://github.com/rrwick/Polypolish) sinning the short reads. The completeness of the draft assembly was evaluated using BUSCO (Manni et al., 2021) and CheckM (Parks et al., 2015), while annotation was carried out using the NCBI Prokaryotic Genome Annotation Pipeline (Tatusova et al., 2016).

### 2.3 Phylogenomic analysis and genome-to-genome metrics

To infer the phylogenetic relationships, a genomic approach was employed. This included all reported genomes for the 16SrX group and 15 representative genomes from other 16Sr groups (Supplementary Material Table S1). To standardize the process, all genomes were re-annotated using Prokka, and identification of orthologous protein clusters was performed using Orthofinder v2.5.2 (https://github.com/davidemms/OrthoFinder). For phylogenetic analyses, alignments of concatenated amino acid sequences of single-copy core genes were constructed using MAFFT v7.505 (https://mafft.cbrc.jp/alignment/software/). Phylogenetic trees were constructed with IQTREE (http://www.iqtree.org/) using an automatic substitution model and ultrafast bootstrap analysis with 1000 replicates. Average nucleotide identity (ANI) was calculated for phytoplasmas from 16SrX group using fastANI v1.1 (https://github.com/ParBLiSS/FastANI).

### 2.4 Characterization of effectors and immunodominant membrane proteins

To identify effector proteins, we utilized well-established methodologies (Bai et al., 2006; Fernández et al., 2019b; Music et al., 2019). Signal peptides were predicted using the SignalP v6.0 server (https://services.healthtech.dtu.dk/services/SignalP-6.0/) and further analyzed with the TMHMM-2.0 server (https://services.healthtech.dtu.dk/service.php?TMHMM-2.0). Proteins that lacked transmembrane helix domains beyond the signal peptide were classified as potential secreted proteins. Finally, the candidate secreted proteins were subjected to reciprocal BLASTp searches (E-value ≤ 1e-05) against proteins from the aster yellows witches’-broom (AYWB) phytoplasma (taxid: 322098) to identify homologous Secreted Aster Proteins (SAPs) (Bai et al., 2006). Identification of immunodominant membrane protein type imp was conducted by BLASTp using the imp sequence of ‘*Ca*. Phytoplasma mali’ AT (accession CAP18237.1) as query. Structural and phylogenetics analysis were made in Geneious R.10 software (Biomatters Ltd., Auckland,New Zealand). The final tree was constructed using the nucleotide sequences of imp gene representing the different imp-profiles based in previous studies (Danet et al., 2011, Bohunická et al., 2018) in IQTREE (http://www.iqtree.org/) using an automatic substitution model and ultrafast bootstrap analysis with 1000 replicates.

## 3. Results and discussion

### 3.1 Assembly and key features of *Ca*. Phytoplasma pyri strain P1 draft genome

*De novo* final assembly consisted of 16 contigs with a total length of 575,431 bp (N50: 88,555 bp), a GC content of 20.35%, and 125X coverage for Illumina reads (Table 1, Figure 1). During the annotation process, a total of 536 genes were identified, including 500 CDSs (with protein), 1 complete rRNA operon (5S, 16S, and 23S), and 30 tRNAs. The functional annotation of CDSs using BlastKOALA (https://www.kegg.jp/blastkoala/) assigned 298 of 500 CDSs (59.6%) to orthologs in the KEEG database. The ‘*Ca*. Phytoplasma pyri’ draft genome contains 170 CDSs assigned to Metabolism, 219 CDSs assigned to Genetic Information Processing, and 39 CDSs assigned to Signaling and Cellular Processes (Figure S1). BUSCO analysis indicated a completeness of 91,40% (mollicutes_abd10), while CheckM reported a completeness of 98.86%. Within the 16SrX group, the complete genome of ‘*Ca*. Phytoplasma mali’ strain AT (16SrXA) (ACCESSION GCA_000026205.1) has been described (Kube et al., 2008), and there are two reports of partial genomes for ‘*Ca*. Phytoplasma prunorum’ (16SrX-B) strains LNS1 (GCA_036924415.1) and ESFY1 (GCA_036924395.1) (Fonseca et al., 2024). The partial genome of ‘*Ca*. Phytoplasma pyri’ strain P1 described here shows completeness values comparable to those of the complete genome of ‘*Ca*. Phytoplasma mali’, demonstrating the quality of the obtained assembly (Table 1). The 16S rRNA gene sequence revealed 100% identity with the reference strain of ‘*Ca*. Phytoplasma pyri’ (AJ542543). Additionally, the *in silico* RFLP profile obtained from the iPhyClassifier server (Zhao et al., 2009) matches the reference pattern for 16Sr group X, subgroup C (GenBank accession: AJ542543), with a similarity coefficient of 1.00. The phytoplasma under study is classified as a member of 16SrX-C.

**Table 1.**
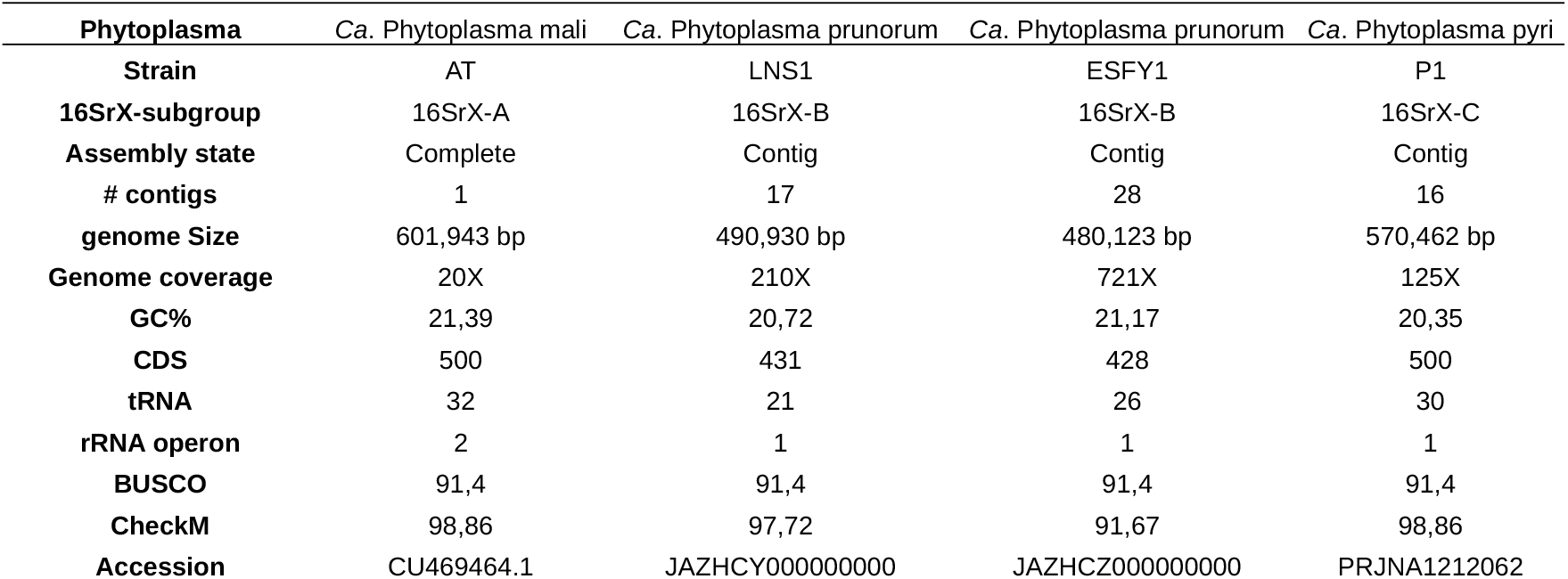
Genomics features of ‘*Ca*. Phytoplasma pyri’ P1 and related genomes from 16SrX group.

**Figure 1.**
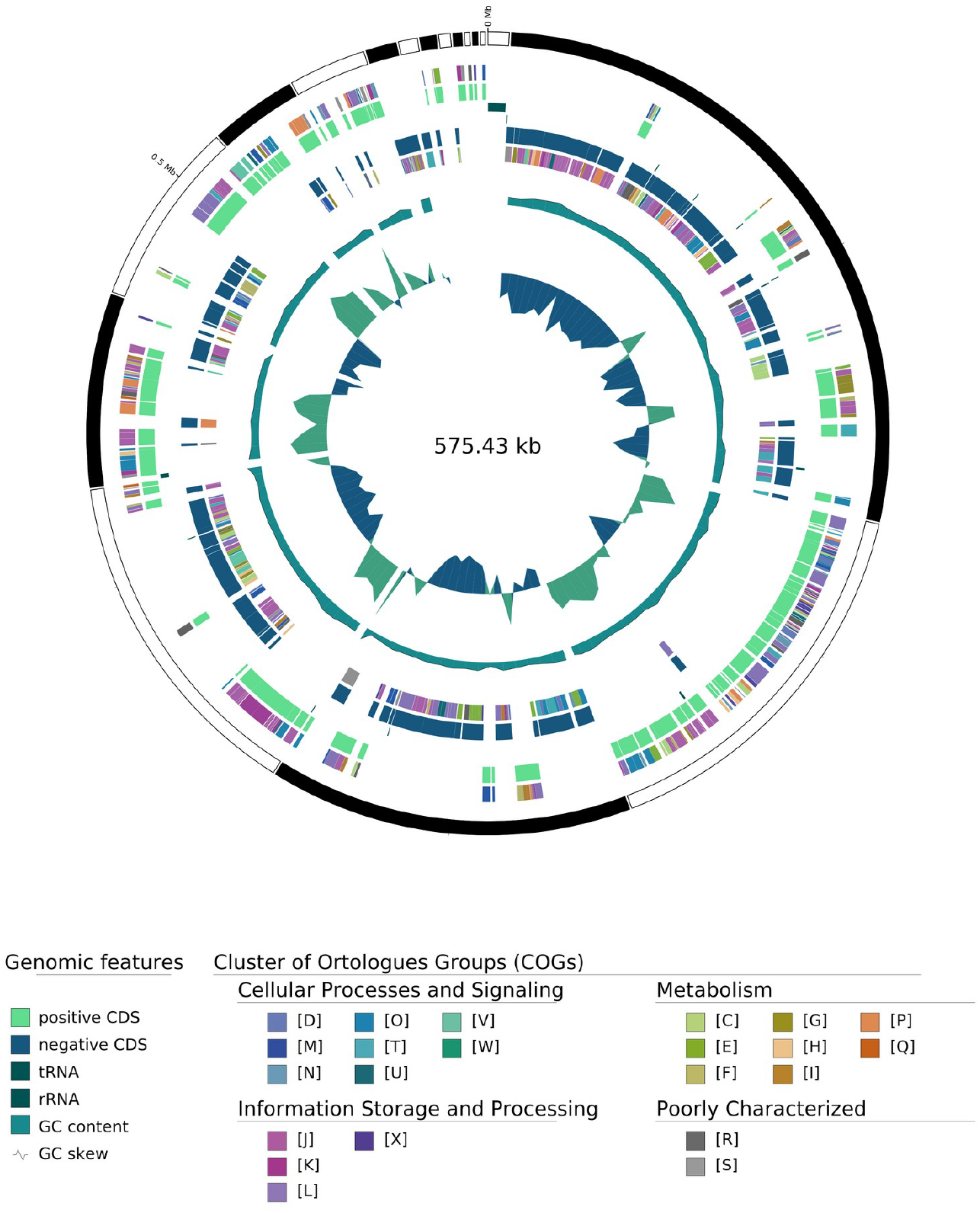
Circular map representation of ‘*Ca*. Phytoplasma. pyri’ strain P1 draft-genome. Labelling from outside toinside: Contigs, COGs on the forward strand, CDS, tRNAs, and RNAs on the foward strand, CDS,tRNAs, and RNAs on the reverse strand, COGs on the reverse strand, GC content and GC skew.

### 3.2 Phylogenomic analyses and genome-to-genome metrics

Comparative genomics based on orthologues identification between draft genomes of ‘*Ca*. Phytoplasma pyri’ strain P1 and representative genome sequence of 18 ‘*Ca*. Phytoplasma species’ (Supplementary Table S1) reveals the presence of 143 single copy genes (CDSs) shared by all the genomes. Phylogenetic reconstruction was undertaken based on concatenated alignment of 46.492 aa (Best-fit model: JTTDCMut+F+I+G4). The resulting phylogenetic tree grouped all representatives of the 16SrX group (‘*Ca*. Phytoplasma mali’, ‘*Ca*. Phytoplasma prunorum’, and ‘*Ca*. Phytoplasma pyri’) within the same clade (Figure 2), separate from the other species, with a high bootstrap value (100), suggesting a relatively recent common ancestor and a close evolutionary relationship between these species. Despite the proximity observed among ‘*Ca*. Phytoplasma pyri’ P1, ‘*Ca*. Phytoplasma mali*’* AT, and the ‘*Ca*. Phytoplasma prunorum’ strains (LNS1 and ESFY1) in the phylogenetic tree, the Average Nucleotide Identity (ANI) analysis provides critical evidence to consider them as distinct species (Figure 3). While the ANI values between these phytoplasmas are relatively high, indicating a recent common ancestor, none of the values exceed the 95–96% threshold typically considered the boundary for bacterial species delineation (Kim et al., 2014, Hugenholtz 2021) and within ‘*Ca*. Phytoplasma species’ (Wei and Zhao, 2022, Bertaccini et al., 2022). Specifically, although the ‘*Ca*. Phytoplasma prunorum’ strains LNS1 and ESFY1 (Fonseca et al., 2024) exhibit near-complete identity (99.6–100%), suggesting they could be classified as the same strain or subspecies, the ANI values between ‘*Ca*. Phytoplasma *pyri’* P1 and the other two species (around 90.3–90.5%), as well as between ‘*Ca*. Phytoplasma mali*’* AT (Kube et al., 2008) and ‘*Ca*. Phytoplasma prunorum’ (around 90.6–90.7%), are consistently below this threshold. The phylogenetic proximity and ANI values below the species threshold strongly supports the classification of ‘*Ca*. Phytoplasma pyri’ P1, *Ca. ‘Ca*. Phytoplasma mali*’*, and ‘*Ca*. Phytoplasma prunorum’ as three distinct, albeit closely related, species. This result supports previous phylogenetical, serological and biological separations of these three Phytoplasma species (Seemüller and Schneider 2004).

**Figure 2.**
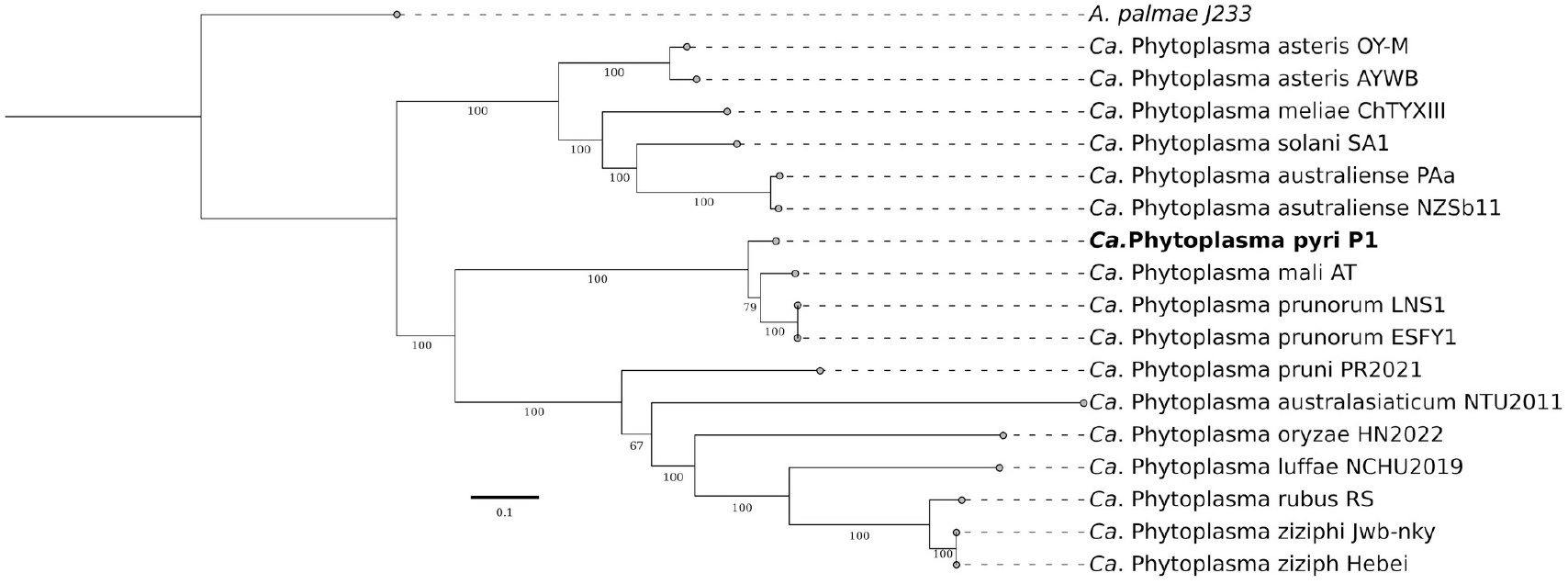
Molecular phylogeny based on amino acid sequences of 143 core genes concatenated. The numbers on branches indicate the level of bootstrap support (1,000 replicates). The scale bar represents the number of substitutions per site. The sequence obtained in this paper is highlighted in bold.

**Figure 3.**
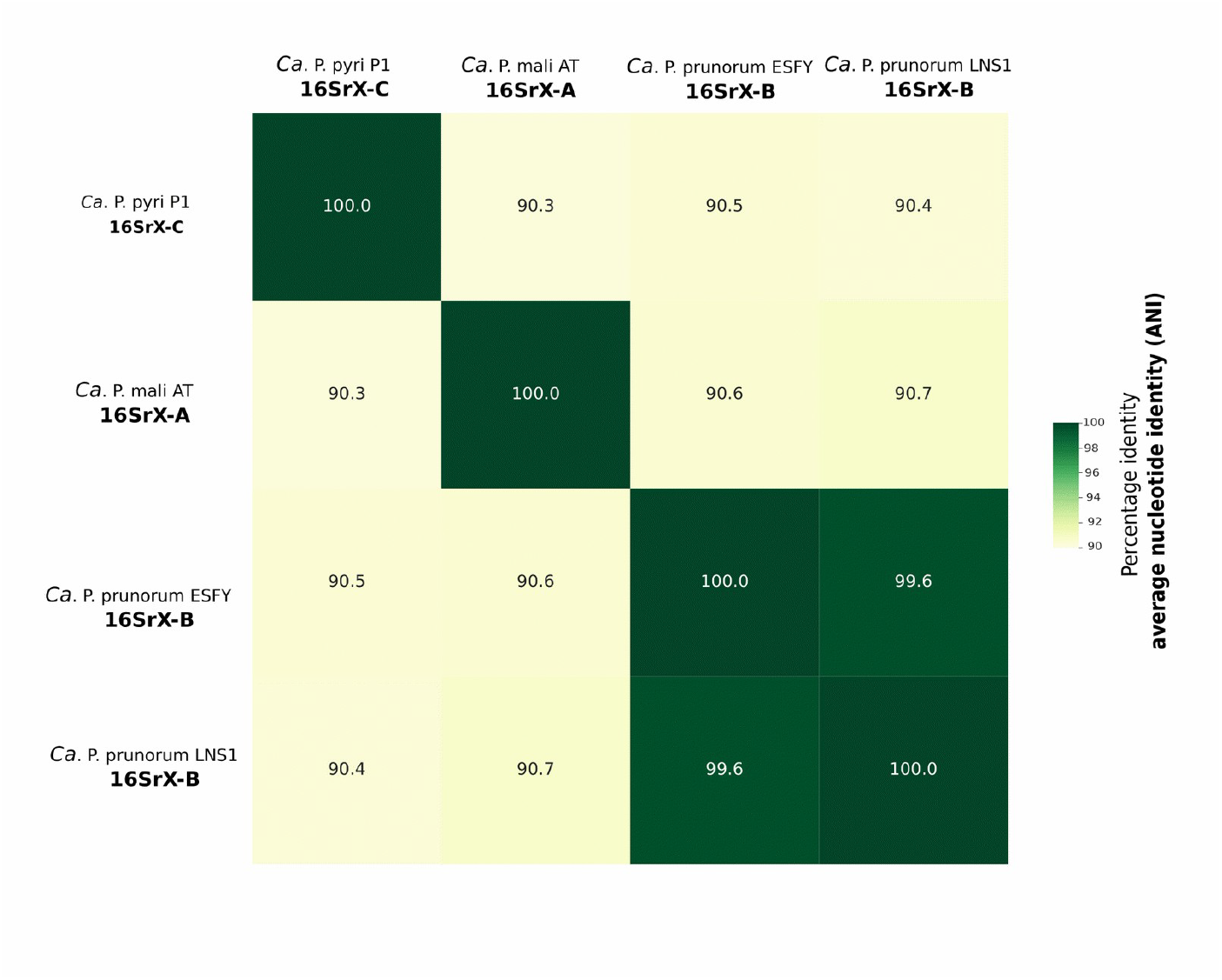
Heatmap generated using fastANI showing the Average Nucleotide Identity (ANI) values (%) among ‘*Ca*. Phytoplasma pyri’ P1 (16SrX-C), ‘*Ca*. Phytoplasma *mali’* AT (16SrX-A), and ‘*Ca*. Phytoplasma *prunorum’* strains ESFY and LNS1 (both 16SrX-B).

### 3.3 Identification of effector proteins

In the search for pathogenic effectors, the presence of 17 secreted proteins (SP positive without transmembrane domain signals after the signal peptide) was identified. Among these 17 proteins, orthologs to the effectors SAP39, SAP41, SAP45, SAP48, and SAP72 were found, but not to the classical effectors SAP05, SAP11, SAP54, and tengu. Additionally, other non-classical effectors, such as the protein PM19_00185 (Strohmayer et al., 2019) or the effector PME2 (Mittelberger et al., 2019), have been described in the phytoplasma ‘*Ca*. Phytoplasma mali’, fulfilling interesting roles in pathogenicity and its interaction with the plant. For the case of the phytoplasma ‘*Ca*. Phytoplasma pyri’ strain P1, we have identified a homolog of the effector PM19_00185, with a size of 210 amino acids, no signal peptide, and 80.70% identity compared to the reference sequence (WP_041186049.1). Currently, it is proposed that HflB proteases and AAA+ ATPases participate in the virulence of the Apple Proliferation phytoplasma, but the specific mechanisms remain unknown (Seemüller et al., 2011; Schneider et al. 2014; Seemüller et al., 2017). In the partial genome of ‘*Ca*. Phytoplasma pyri’ we have identified 6 proteins related to these, with levels of conservation, which may indicate a conserved mechanism in phytoplasmas of the 16SrX group. However, more experimental evidence is needed to support this hypothesis. The presence of fragmented genomes may be influencing the ability to identify all effectors. It is necessary to continue generating high-quality genomic resources to ensure adequate representativeness.

### 3.4 Characterization of immunodominant membrane protein type imp

We have identified a homologue of the Imp immunodominant membrane protein in the ‘*C*a. Phytoplasma pyri’ draft genome. The organization of Imp surrounding genes (pyrG-imp-dnaD-rnc) was conserved (Figure 4.A). The deduced Imp protein sequence comprises 168 amino acids, with a predicted transmembrane helix region spanning residues 9-29 and extracellular hydrophilic domain in residues 30-168. Same organization was described for Imp encoding protein in strains LNS1/ESFY and AT (Figure 4.B). We have also found that the amino acid identity between the Imp described for the P1 isolate was 47.65% compared to the sequence of the EFSY1 and LNS1 isolates, while it was 49.71% compared to the AT isolate.

**Figure 4.**
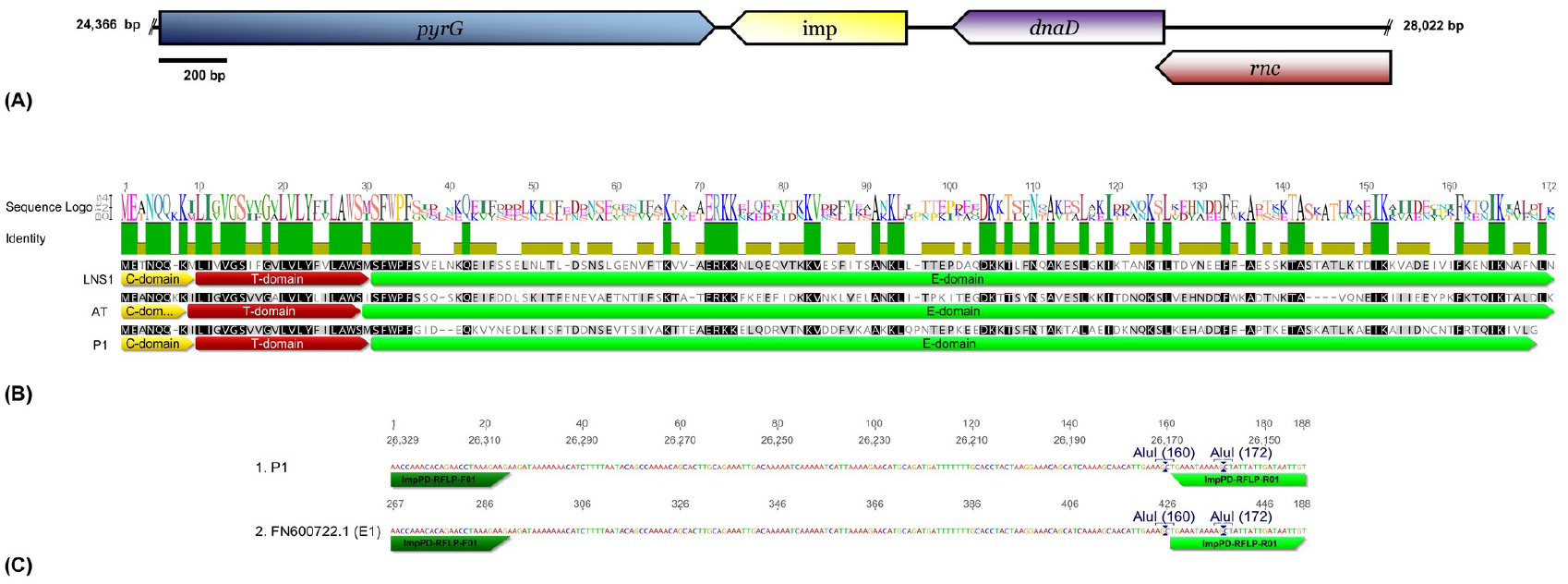
Molecular characterization of the *imp* encoding gene in ‘*Ca*. Phytoplasma pyri’ strain P1. (A) Gene organization of the *imp* genomic context. (B) MSA alignment of the *imp* amino acid sequences from LNS1, AT, and P1 strains (C-domain: internal domain, T-domain: helix domain, E-domain: external hydrophilic domain). (C) Schematic representation of the nucleotide sequence amplified with the ImpPD-RFLP-F01/ImpPD-RFLP-R01 primers (Bohunická et al., 2018) showing *AluI* restriction sites in the P1 sequence and a representative sequence of the E1 *imp* genotype.

The imp gene was used as molecular marker to describe the diversity in phytoplasmas form 16SrX-group (Dannet et al., 2011, Bohunická et al., 2018) showing shows high inter- and intraspecies variability and demonstrating the existence of inter-species recombination between ‘*Ca*. Phytoplasma pyri’ and ‘*Ca*. Phytoplasma prunorum’ species. To infer the phylogenetic relationships of ‘*Ca*. Phytoplasma pyri’ strain P1 based on imp sequence a phylogenetic tree was constructed using representative sequences of different described genotypes (Bohunická et al.,2018). The final tree grouped the strain P1 within the sequences of representative genotype E1/I15 (Figure 5) in the same cluster. This grouping is consistent with the results of the *in silico* RFLP profile of the fragment amplified with the primers ImpPD-RFLP-F01/ImpPD-RFLP-R01 (Bohunická et al., 2018) and the digestion of this fragment with the *Alu*I enzyme (Figure 4.C). These results confirm that the sequenced isolate corresponds to the previously described imp genotype E1.

**Figure 5.**
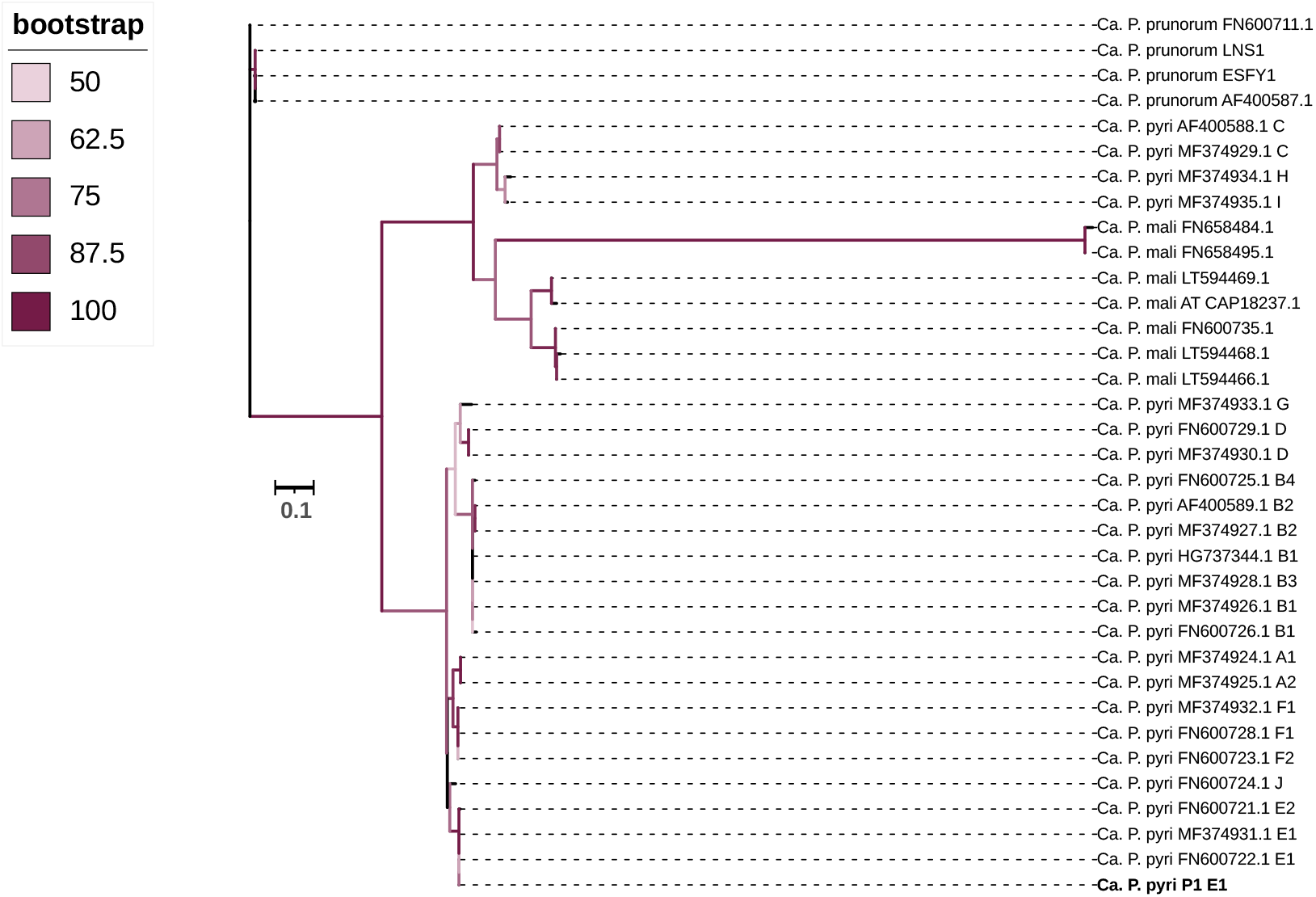
Phylogenetic tree of ‘*Ca*. Phytoplasma species’ reconstructed using the Maximum Likelihood (ML) method under the TVM+F+G4 model. The tree includes sequences from ‘*Ca*. Phytoplasma pyri’, ‘*Ca*. Phytoplasma mali’, and ‘*Ca*. Phytoplasma prunorum’, with GenBank accession numbers indicated. Bootstrap values (percentages) are shown at branch nodes, with values below 50% omitted for clarity. The scale bar represents 0.1 substitutions per site. For ‘*Ca*. Phytoplasma pyti’ sequences, the inferred genotype based on restriction analysis of the imp gene (Bohunická et al., 2018) is provided after the GenBank accession number.

Imp displays significant genetic variability within phytoplasmas and undergoes strong positive selection pressure, indicating its role in interactions with the environment and the host (Kakizawa et al., 2009). Studies have shown that Imp binds not only to plant actin (Boonrod et al., 2012) butalso to various insect proteins, establishing a correlation between Imp binding and vector status (Siampour et al., 2011; Trivellone et al., 2019). Since Imp likely plays a crucial role in host–pathogen interactions, characterizing its diversity in ‘*Ca*. Phytoplasma pyri’ could provide valuable nsights into the epidemiology, symptom development, and other facets of pear decline disease.

## 4. Conclusions

Advances in genomics have provided critical insights into their genetic diversity and interactions with hosts, supporting the development of strategies for managing phytoplasma-associated diseases. This study presents the draft genome of *‘Ca*. Phytoplasma pyri’ strain P1, offering insights into its genetic composition and pathogenic potential. We identified 536 genes in the genome, including those involved in metabolism, genetic information processing, and signaling, which are commonly found in phytoplasmas. The phylogenetic analysis places *‘Ca*. Phytoplasma pyri’ strain P1 within the 16SrX group, supporting its classification as a distinct species from *‘Ca*. Phytoplasma mali’ and ‘*Ca*. Phytoplasma prunorum’, despite their close evolutionary relationships. In our search for pathogenic effectors, 17 secreted proteins were identified, some of which are homologs of known effectors such as SAP39 and SAP41, and others fulfilling roles in pathogenicity similar to those described in related phytoplasmas. We also identified six conserved proteins related to HflB proteases and AAA+ ATPases, which may be involved in virulence mechanisms. Additionally, a homologue of the Imp immunodominant membrane protein was identified, showing a conserved gene organization across strains. Although some of the identified proteins, such as PME2, were not homologous in the *‘Ca*. Phytoplasma pyri’ draft genome. This study underscores the importance of continued genomic exploration to provide a comprehensive understanding of phytoplasma species and their impact on plants. Furthermore, it represents the first global report of a *‘Ca*. Phytoplasma pyri’ genome, marking a significant advancement in the global phytoplasma genomics landscape and paving the way for future research in this area.

## Data availability

All data generated or analyzed during this study are included in this published article. Accession Numbers: The whole-genome sequence of *Candidatus* Phytoplasma pyri strain P1 can be accessed via National Center for Biotechnology Information (NCBI) database with PRJNA1212062 Bioproject number.

## Supplementary Materials

**Supplementary Figure S1:**
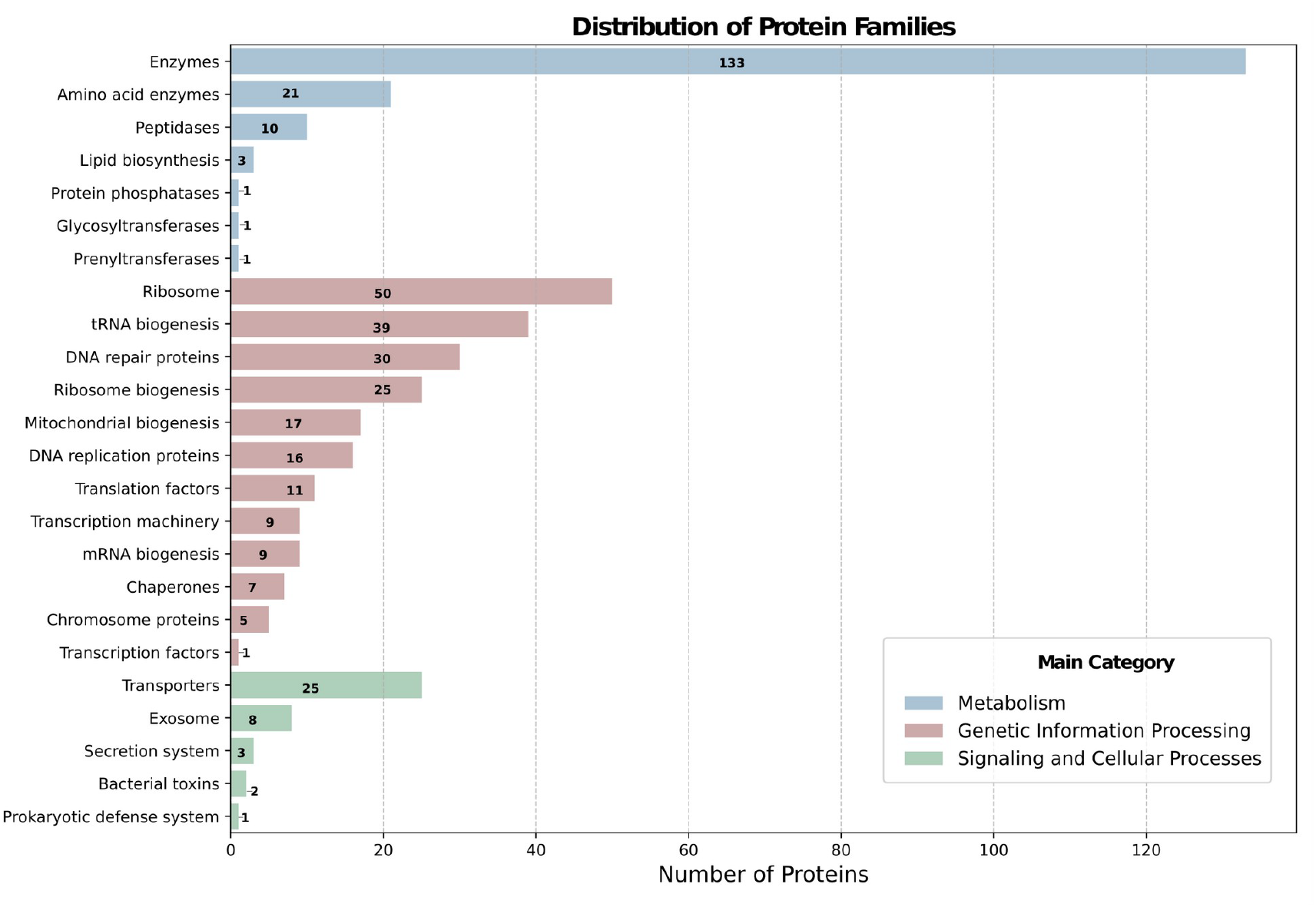
List of protein families associated to KEEG in the ‘*Ca*. Phytoplasma pyri’ draft genome

